# An increase of *NPY1* expression leads to inhibitory phosphorylation of PIN-FORMED (PIN) proteins and suppression of *pinoid* (*pid*) null mutants

**DOI:** 10.1101/2025.07.14.664678

**Authors:** Michael Mudgett, Zhouxin Shen, Ruofan Kang, Xinhua Dai, Steve Briggs, Yunde Zhao

## Abstract

The PINOID (PID) protein kinase is required for flower initiation in Arabidopsis. The *pid* mutants fail to initiate flowers on inflorescences, a phenotype that is mimicked by disrupting either the *NAKED PINS IN YUC MUTANTS* (*NPY)* gene family or *PIN FORMED 1* (*PIN1*). Both PID and NPY1 have been reported to positively modulate PIN-mediated polar auxin transport. Here, we show that overexpression of *NPY1* (*NPY1 OE*) completely suppressed *pid* null mutants, demonstrating that *NPY1* functions downstream of *PID*. *NPY1 OE* triggered phosphorylation of PIN proteins at multiple sites that are mostly different from the previously characterized phosphorylation sites regardless of the presence of *PID*. Phosphorylation of the newly identified PIN sites in *NPY1 OE* plants likely leads to the inhibition of PIN functions, as we previously showed that *pid* is suppressed by decreasing *PIN1* gene dosage or decreasing PIN1 activity. Furthermore, we show that the Ser/Thr rich C-terminal motif in NPY1 is phosphorylated and is required for *pid* suppression by *NPY1 OE.* Overexpression of *NPY1* that lacked the C-terminal motif (*NPY1ΔC*) failed to rescue *pid*, but overexpression of *NPY1ΔC* was still able to trigger phosphorylation of PIN proteins including PIN2, which is known to cause agravitropic roots when mutated. *NPY1ΔC* overexpression plants displayed a complete loss of root gravitropic response, likely caused by PIN2 phosphorylation. Our results suggest a pathway for auxin mediated-flower initiation, in which PID regulates NPY1 accumulation and/or activity, and subsequently, NPY1 triggers phosphorylation of PIN proteins and inhibition of PIN functions.

## Introduction

Auxin is required for flower initiation in Arabidopsis and other plants (Cheng and Zhao, 2007; Yamaguchi et al., 2013). Several Arabidopsis mutants and mutant combinations fail to initiate flowers on inflorescences, despite the fact that they can undergo the normal transition from vegetative growth to reproductive growth, resulting in the formation of pin-like inflorescences (Bennett et al., 1995; Przemeck et al., 1996; Galweiler et al., 1998; Cheng et al., 2007). All of the known pin-like Arabidopsis mutants are caused by defects in some aspects of auxin biology. *PIN-FORMED 1* (*PIN1*) was the first characterized gene that causes the formation of pin-like inflorescences when compromised (Galweiler et al., 1998). PIN1 is an auxin efflux carrier that transports auxin from the cytosol into the extracellular space (Yang et al., 2022). Plants grown on media containing the polar auxin transport inhibitor N-1-naphthylphthalamic acid (NPA) phenocopy *pin1* mutants (Okada et al., 1991). It has been shown that NPA can directly bind to PIN1 to inhibit auxin transport (Yang et al., 2022). Disruption of *PINOID* (*PID*), which encodes a Ser/Thr protein kinase, also leads to the formation of pin-like inflorescences (Smyth and Alvarez, 1995; Christensen et al., 2000). Overexpression of *PID* leads to phenotypes similar to the well-characterized auxin resistant mutants, linking *PID* functions to auxin (Christensen et al., 2000). *NAKED PINS IN YUC MUTANTS 1*(*NPY1)* (Cheng et al., 2007), which is also called *ENHANCER OF PINOID* (*ENP*) (Treml et al., 2005) and *MACCHI-BOU 4* (*MAB4*) (Furutani et al., 2007), was isolated from genetic screens for *pid* enhancers and for enhancers of *yuc1 yuc4* double mutants (Cheng et al., 2006), which are defective in auxin biosynthesis (Cheng et al., 2007). The *yuc1 yuc4 npy1* triple mutants produce pin-like inflorescences similar to those in *pin1* and *pid* (Cheng et al., 2007). Inactivation of *NPY1* and its close homologs *NPY3* and *NPY5* also causes the failure of floral initiation from inflorescences (Cheng et al., 2008). The *yuc1 yuc2 yuc4 yuc6* quadruple mutants also develop small pin-like structures (Cheng et al., 2006).

Although the aforementioned genes are known to participate in auxin-mediated flower development, the mechanisms by which the genes regulate floral development and the relationships between them are not fully understood. The predominant model regarding the relationship among *PIN1*, *PID*, and *NPY* genes is centered on regulating PIN polarity, localization, and activity by PID and NPY proteins. It was reported that PID directly phosphorylates the hydrophilic loop of PIN proteins, resulting in changes in PIN polarity and activation of PIN-mediated auxin export (Friml et al., 2004; Michniewicz et al., 2007). NPYs were suggested to regulate PIN internalization through the endocytosis pathway (Furutani et al., 2011). Recently, it was reported that NPY1, PID, and PIN1 form a protein complex at the plasma membrane and that the recruitment of NPYs to the plasma-membrane by PIN limits the lateral diffusion of PINs, thus maintaining PIN polarity (Glanc et al., 2021). Phosphorylation of PIN by PID enhances the recruitment of NPY to plasma membrane, which then promotes PIN phosphorylation by recruiting or interacting with AGC kinases, thus forming a self-reinforcing loop to maintain PIN polarity (Glanc et al., 2021). Other protein kinases including D6 PROTEIN KINASE are also involved in phosphorylating PIN proteins and activating PIN-mediated auxin transport (Willige et al., 2013; Barbosa et al., 2014).

Recently, we discovered that *pid* null mutant phenotypes were suppressed when one copy of the *PIN1* gene is inactivated (heterozygous *pin1* mutants) (Mudgett et al., 2023). The *pid* null mutants were also suppressed by the PIN1-GFPHDR fusion in which *GFP* was precisely inserted into the *PIN1* gene via CRISPR/Cas9-mediated homologous recombination (Mudgett et al., 2023). PIN1-GFPHDR is less active than wild type (WT) PIN1, indicating that *pid* is suppressed by lowering either PIN1 activity or *PIN1* gene dosage. This new type of genetic interaction between *pid* and *pin1,* called haplo-complementation, renders *PID* unnecessary for flower development when expressing only 50% of *PIN1*, whereas the presence of 0% or 100% *PIN1* makes *PID* essential. The observed genetic interaction between *pid* and *pin1* suggests a much more complex relationship between the two genes than the current model of PIN1 phosphorylation by PID directly (Mudgett et al., 2023). Moreover, it was known that *pid* mutants produce more cotyledons and true leaves than wild type plants whereas *pin1* mutants make fewer cotyledons and fewer true leaves compared to wild type plants (Bennett et al., 1995). The opposite cotyledon/true leaf phenotypes of *pid* and *pin1* cannot be accounted for by PID-mediated direct phosphorylation/activation of PIN1.

In this study, we analyzed the dosage effects of *NPY1* on *pid* phenotypes. We show that overexpression of *NPY1* (*NPY1 OE*) completely suppressed *pid* null mutants, demonstrating that *NPY1* and *PIN1* have opposite dosage effects on *pid* mutants. Furthermore, we find that *NPY1 OE* led to an increase of PIN phosphorylation, suggesting that *NPY1 OE* -triggered PIN phosphorylation actual inhibits PIN functions. Moreover, we discovered that the C-terminal motif of NPY1, which contains 30 amino acid residues with 50% Ser/Thr, is required for suppression of *pid* by *NPY1 OE*. Overexpression of *NPY1* that lacked the C-terminal motif (*NPY1ΔC*) did not suppress *pid*, but caused auxin resistance and a complete loss of gravitropism. Overexpression of *NPY1ΔC* also increased phosphorylation of PIN proteins including PIN2, which causes agravitropic roots when mutated, further suggesting that phosphorylation of certain residues/regions of PIN proteins inhibits PIN functions. Our results demonstrated that NPY1 functions downstream of PID during flower development. In addition, our results indicate that NPY1 affects PIN phosphorylation, which inhibits PIN function and which may account for the suppression of *pid* by *NPY1 OE*.

## Results

### Protein null pin1 mutants further confirm haplo-complementation of pid mutants by pin1

We previously showed that *pid* null mutants were rescued by either PIN1-GFPHDR fusion or the presence of only one copy of functional *PIN1* gene, suggesting that a reduction of PIN1 activity or *PIN1* gene dosage is sufficient to restore the fertility of *pid* (Mudgett et al., 2023). Because the *pin1* mutants in our previous study had the potential to produce truncated PIN1 proteins, we could not completely rule out the possibility that the predicted truncated PIN1 proteins might have played a role in rescuing *pid* (Mudgett et al., 2023). To further clarify the mechanisms of rescuing *pid* by heterozygous *pin1* mutants, we used CRIPSR/Cas9 to delete the coding region of *PIN1* so that no truncated PIN1 proteins would be produced (Figure 1A). We obtained two *pin1* mutants, which lacked almost the entire coding region of *PIN1* and which were unlikely to produce any truncated PIN1 proteins (Figure 1A). The two *pin1* mutants displayed strong pin-like phenotypes (Figure 1B & 1C). In the heterozygous *pin1* mutant backgrounds, the *pid* null mutants no longer produced *pin*-like inflorescences and were able to set seeds (Figure 1B & 1C). Rescuing *pid* by heterozygous *pin1* mutants was not specific to a particular *pid* allele. Both T-DNA insertion (Figure 1B) and CRISPR deletion *pid* mutants (Figure 1C), which were previously described as *pid* null mutants (Mudgett et al., 2023), were rescued by the heterozygous *pin1* full deletion mutants (Figure 1), demonstrating that *pid* mutants were rescued by a reduction of *PIN1* gene dosage and that suppression of *pid* mutant phenotypes by heterozygous *pin1* mutants in our previous study was not caused by the predicted truncated PIN1 proteins.

**Figure 1.**
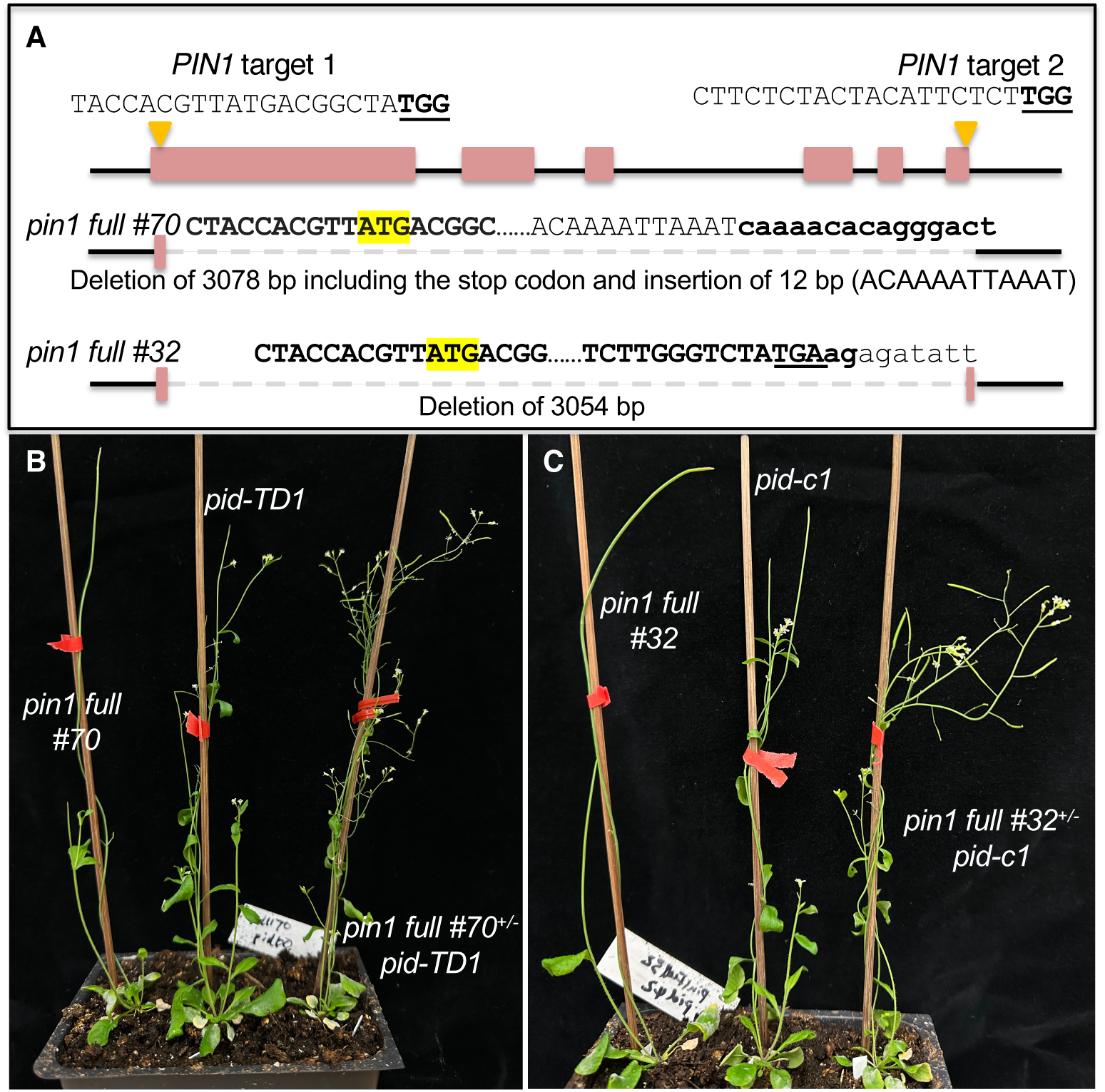
A complete deletion of one copy of the *PIN1* gene rescues *pid* null mutants. A) *PIN1* gene is deleted using CRISPR/Cas9 gene editing technology. The two guide RNA target sites are shown with PAM underlined and in bold. The PIN1 start codon ATG is highlighted in yellow. The sequences flanking the deletions are shown. The two mutants have less than 18 bp of PIN1 coding sequence and are unlikely to produce any PIN1 proteins. B) Heterozygous *pin1 full #70,* which had a deletion of 3078 bp of the *PIN1* gene, restored the fertility of *pid-TD1*, a T-DNA insertion mutant that is completely sterile in WT *PIN1* background. C) *pin1 full #32*, when heterozygous, rescued *pid-c1*, a null *pid* allele.

### NPY1 and PIN1 have different effects on pid mutants - a reduction of NPY1 gene dosage cannot rescue pid

Inactivation of *NPY1* and its close homologs *NPY3* and *NPY5* leads to the development of pin-like inflorescences (Cheng et al., 2008). Moreover, *npy1 pid* double mutants fail to develop any cotyledons, a phenotype that was also observed in *pin1 pid* double mutants (Furutani et al., 2004; Treml et al., 2005; Furutani et al., 2007; Cheng et al., 2008). Therefore, we hypothesized that a reduction of *NPY1* gene dosage might also be able to rescue *pid* mutants. We genotyped F2 plants from a cross of *pid-TD1* and *npy1-2* to identify plants with either *npy1-2 pid-TD1*^+/-^ or *npy1-2*^+/-^ *pid-TD1* genotype to investigate the effects of gene dosage on genetic interactions between the two mutants. It was very clear that heterozygous *npy1-2* did not rescue *pid-TD1* (Supplemental Figure 1). Interestingly, heterozygous *pid-TD1* further enhanced *npy1-2*, which is fertile and which does not produce pin-like structures (Supplemental Figure 1). Plants with the *npy1-2 pid-TD1*^+/-^ genotype were essentially sterile and produced pin-like structures (Supplemental Figure 1). Our results suggest that gene dosage effects of *NPY1* and *PIN1* on *pid* are different.

### Dosage effects of NPY1 and PIN1 on pid are opposite - Increased NPY1 gene expression is sufficient to suppress pid null mutants

We placed *NPY1* gene under the control of the *UBIQUITIN 10* promoter to strongly express *NPY1* (named as *NPY1 OE)*. We directly transformed the construct into a *pid-c1* segregation population through Agrobacterium-mediated transformation. Among the 84 T1 plants we genotyped, we obtained 15 homozygous *pid-c1* mutants. Interestingly, none of the *pid-c1* plants produced any pin-like inflorescences and all were fully fertile, demonstrating that increases of *NPY1* gene expression were sufficient for the suppression of *pid-c1*. We genotyped T2 progenies from two *pid-c1* heterozygous T1 plants (#68 and # 83) for the presence of *pid-c1* and for *pid-c1* zygosity. We used mCherry signal, which was included in the *NPY1 OE* construct, as a proxy to determine the presence and absence of the *NPY1* transgene. For each line, we identified T2 plants without the *NPY1* transgene and without the *pid-c1* mutation (called WT-68 and WT-83, respectively). We also isolated T2 plants that contained the *NPY1* overexpression construct, but did not have the *pid-c1* mutation (called *NPY1 OE* #68 in WT, and *NPY1 OE* #83 in WT). Finally, we identified T2 plants that were *pid-c1* homozygous and that had the *NPY1* transgene (called *NPY1 OE #68* in *pid-c1* and *NPY1 OE #83* in *pid-c1*). These genetic materials enabled us to compare the same *NPY1 OE* transgenic event in different genetic backgrounds.

As shown in Figure 2A and Supplemental Figure 2, both *NPY1 OE #68* and *#83* lines rescued *pid-c1,* which lacks almost the entire *PID* coding region and which is completely sterile under our laboratory conditions (Mudgett et al., 2023). Overexpression of *NPY1* in *pid-c1* completely eliminated the development of pin-like inflorescences and led to the production of plenty of fertile siliques (Figure 2A & Supplemental Figure 2). *NPY1 OE* lines had notable developmental phenotypes: *NPY1 OE* plants had reduced petiole length and had shorter stature compared to WT plants (Supplemental Figure 2). At the young adult stage, *NPY1 OE* in WT and in *pid-c1* appeared very similar, suggesting that overexpression of *NPY1* made *PID* unnecessary for plant development (Supplemental Figure 2). *NPY1 OE* lines, regardless of the presence or absence of the *pid-c1* mutation, developed plenty of flowers with normal appearance (Supplemental Figure 2). Most flowers from *NPY1 OE* in *pid-c1* plants were normal, but occasionally, some flowers had minor defects such as missing a stamen, or developing fused petals with a stamen-like structure (Supplemental Figure 2). In comparison to the suppression of *pid* null mutants by PIN1-GFPHDR or heterozygous *pin1,* suppression of *pid-c1* by *NPY1 OE* appeared to be more complete (Mudgett et al., 2023).

**Figure 2.**
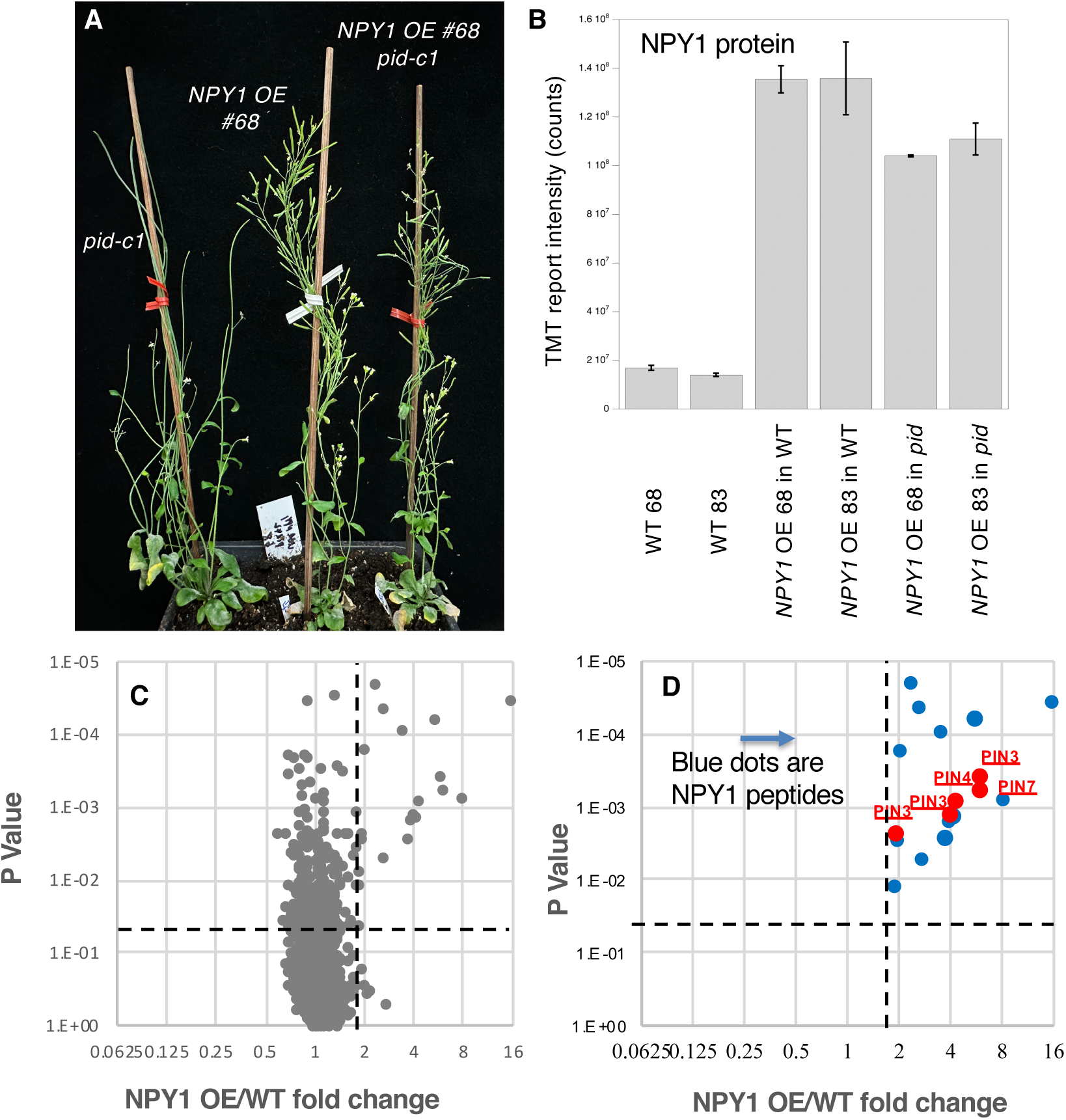
Overexpression of *NPY1* rescues *pid* null mutants and triggers phosphorylation of PIN proteins. A) The *NPY1* overexpression (OE) line #68 rescues the *pid-c1*, a null allele. Note that *pid-c1* makes pin-like inflorescences and is completely sterile. *pid-c1* no longer makes pin-like structures and is completely fertile when NPY1 protein levels are increased. A second *NPY1* overexpression line (#83) also restored the fertility of *pid-c1* (Supplemental Figure S2). B) NPY1 protein levels in the *NPY1 OE* lines in WT background and in *pid* mutants. In line #68, NPY1 protein level is 8.0-fold higher than WT while line #83 has 9.7-fold higher NPY1 protein level than WT. In the absence of *PID*, overexpression of *NPY1* leads to slightly lower NPY1 protein levels than the same transgenic event in WT background. C) Overexpression of *NPY1* leads to an increase of phospho-peptides. The volcano plot shows the fold changes (Log2 scale) of phospho-peptide levels in *NPY1 OE* lines over WT. Data above the horizonal dotted line are statistically significant. The vertical dotted line marks 2-fold change. D) Overexpression of *NPY1* leads to phosphorylation of PIN proteins. Among the phospho-peptides that displayed the most differences between *NPY1 OE* lines and WT are peptides from NPY1 and PIN proteins. In fact, in the right up-Quadrant in C, the only peptides were from NPY1 and PIN proteins.

We also transformed the *NPY1 OE* construct into *pid-TD1* and *pid-TD2*, which are T-DNA insertional null alleles (Mudgett et al., 2023). Both *pid* T-DNA mutants were rescued by overexpression of *NPY1*, demonstrating that *NPY1 OE* could rescue different types of *pid* mutants (Supplemental Figure 3). Some of the siliques in the rescued *pid-TD* plants had only one valve, a phenotype that was likely caused by a partial suppression of *pid-TD* (Supplemental Figure 3). Our results demonstrated that overexpression of *NPY1* rescued *pid* mutants and the suppression of *pid* by *NPY1 OE* was not caused by any background mutations, a particular T-DNA insertion site, or dependence on special *pid* mutations. Our results clearly showed that *NPY1* and *PIN1* had opposite dosage effects on *pid* mutants.

### NPY1 protein accumulates less in the absence of PID

We conducted proteomic analysis to determine the expression levels of *NPY1* in the transgenic lines (Figure 2B). In *NPY1 OE* line #68 in WT, NPY1 protein level increased 8.0-fold compared to non-transgenic WT-68 (Figure 2B). The NPY1 protein level was slightly higher in line #83 than in line #68 (9.7-fold vs 8.0-fold) (Figure 2B). The same *NPY1* transgenic events also led to elevated NPY1 protein levels in *pid-c1* compared to non-transgenic WT (Figure 2B). Interestingly, NPY1 protein levels in the *NPY1 OE* in *pid-c1* plants were significantly lower than *NPY1 OE* in WT. For line #68, NPY1 level was 23% lower in *pid-c1* than in *NPY1 OE* in WT. In line #83, NPY1 protein level was 18.5% lower in *pid-c1* than in WT. Our results suggest that PID likely plays a role in regulating NPY1 protein accumulation.

### NPY1 is phosphorylated and the phosphorylation of NPY1 does not require PID

We identified 15 phospho-peptide of NPY1 (Supplemental Table 1). Among the phosphorylation sites identified, most of them were located in the peptide ANHSPVASVAASSHSPVEK, which has five serine residues and which is located immediately downstream of the conserved NPH3 domain (Supplemental Figure 4). Previous studies have identified two phosphorylation sites in NPY1: S514 and S553 (Matthes et al., 2024). We also detected that S553 was phosphorylated (Supplemental Table 1 and Supplemental Figure 4). Moreover, we identified a couple of phospho-serine residues outside of the C-terminal domain (Supplemental Figure 4). Interestingly, S181 is located in a highly conserved WSYT motif and is located in the region between the BTB domain and the NPH3 domain (Supplemental Figure 4). The S181 residue is conserved among all NPY proteins, but in NPH3, the serine residue is replaced with an alanine (Supplemental Figure 4).

In the absence of PID, all of the 15 phospho-peptide of NPY1 were still phosphorylated (Supplemental Table 1). Quantitatively, all of the peptides except LHEASVK had lower phosphorylation levels in *NPY1 OE* in *pid* plants than in *NPY1 OE* in WT plants. But given that in *pid*, NPY1 protein concentrations were about 20% lower than in WT (Figure 2B), the observed differences in phosphorylation levels could be caused by a decrease of NPY1 protein concentration in *pid*.

### Overexpression of NPY1 leads to increases in phosphorylation of PIN proteins

Overexpression of *NPY1* caused significant changes in the phospho-proteome (Figure 2C). Among the phospho-peptides that were highly enriched in *NPY1 OE* lines compared to WT, the majority of them were NPY1 peptides (Figure 2D). Interestingly, the next highly enriched peptides in *NPY1 OE* lines were from PIN proteins (Figure 2D).

The phospho-peptides AGLQVDNGANEQVGKsDQGGAK from PIN7 and AGLNVFGGAPDNDQGGRsDQGAK from PIN3 were about 6-fold higher in *NPY1 OE* lines than in WT (Table 1). The two peptides mapped to the same region of the two PIN proteins (Supplemental Figure 5). Phospho-peptides in PIN1 and PIN4 were also up-regulated in *NPY1 OE* lines compared to WT (Table 1). We did not detect phospho-peptides from PIN2, probably because PIN2 is not expressed in flowers. The two phosphorylated sites in PIN1 (S271 and S282) are conserved within the PIN family and have been previously identified and characterized (Supplemental Figure 5). Two of the PIN7 phosphorylation sites were highly conserved while the majority of the phosphorylation sites in PIN3, PIN4, and PIN7 were located in the less conserved regions of their respective hydrophilic loops (Supplemental Figure 5).

**Table 1.**
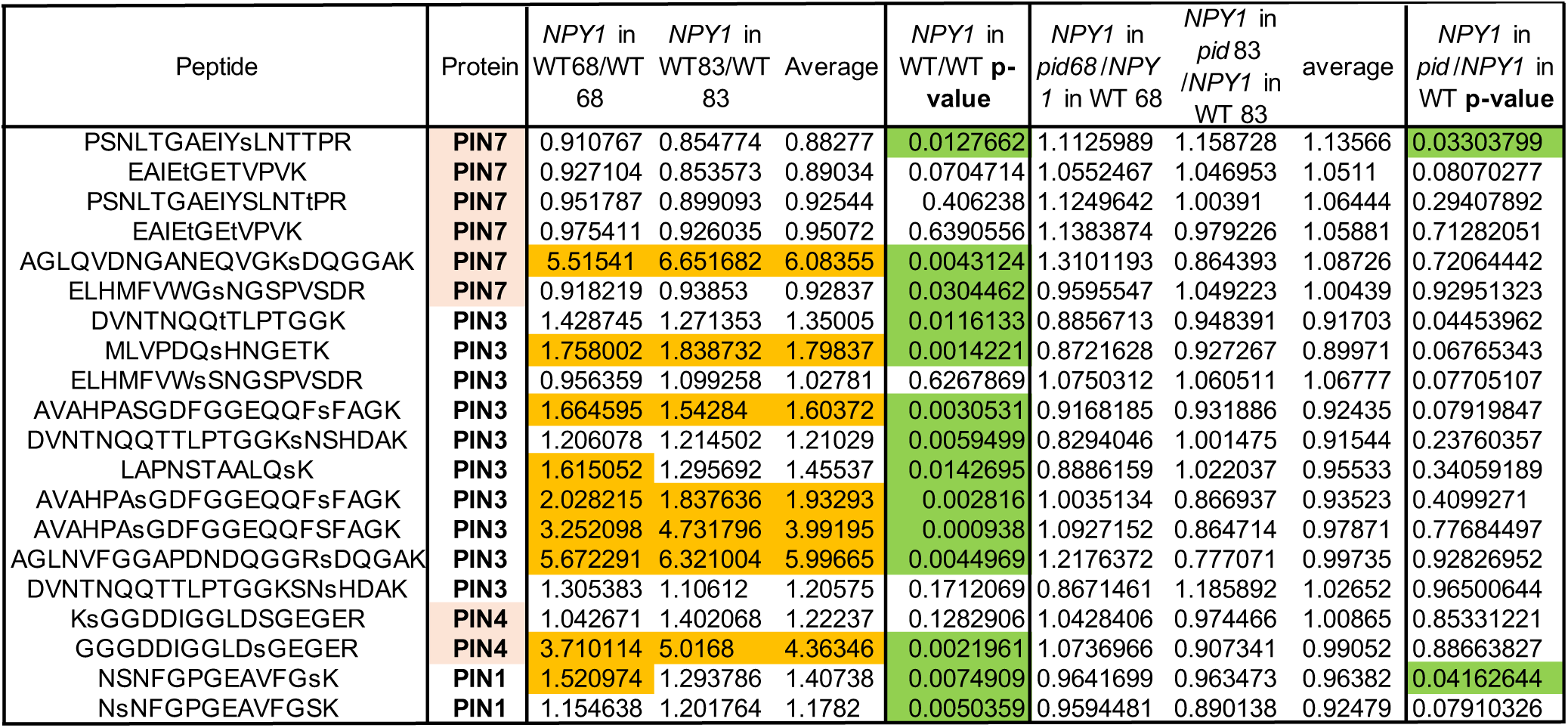
Overexpression of *NPY1* leads to increases of phosphorylation of PIN proteins. *NPY1* in WT 68 refers to *NPY1* overexpression (OE) line #68 in WT background while WT68 refers to plants without the *NPY1 OE* construct segregated from the *NPY1 OE* line #68. *NPY1* in *pid*68 refers to the *NPY1 OE* line #68 in the *pid-c1* homozygous background. The comparisons were made on basis of the same transgenic event of integrating the *NPY1 OE* construct into Arabidopsis genome. Similar nomenclature is used for line #83. Fold change over 1.5 is highlighted in orange and P-value less than 0.05 is highlighted in green.

The absence of *PID* hardly affected PIN phosphorylation triggered by *NPY1 OE* (Table 1), which was consistent with our previous analysis that phosphorylation levels of PIN proteins in *pid* PIN1-GFPHDR plants were not affected compared to WT plants (Mudgett et al., 2023). We noticed that previous studies uncovered four highly conserved phosphorylation sites in PIN proteins, which are named as S1, S2, S3, and S4 sites and which correspond to S231, S252, S290, and S271 in PIN1, respectively (Barbosa et al., 2018) (Supplemental Figure 5). Phosphorylation of S1, S2, S3, or S4 leads to activation of PIN auxin transport and changes in PIN polarity (Lanassa Bassukas et al., 2022). Given that *NPY1 OE* suppressed *pid* mutants and that decreases in *PIN1* gene dosage and PIN1-GFPHDR suppressed *pid*, we concluded that phosphorylation at the newly identified sites in PIN proteins leads to the inhibition of PIN activities.

### The NPY1 C-terminal domain is required for the suppression pid mutants

The NPY1 C-terminal domain is Ser/Thr rich (Supplemental Figure 6), suggesting that the region might be important for regulating NPY1 activity and/or interactions between NPY1 and its partners. The NPY1 phospho-peptides we have identified were largely concentrated in this domain (Supplemental Figure 4). We overexpressed a truncated NPY1 that lacked the 30 C-terminal amino acid residues (called NPY1ΔC), of which 50% are Ser/Thr (Supplemental Figure 6), to determine whether the C-terminal tail of NPY1 was important for NPY1 functions and whether overexpression of *NPY1ΔC* was sufficient for suppression of *pid* mutants. We transformed the *NPY1ΔC* construct into a *pid-TD1* segregating population. Among the T1 plants that were identified to be *pid-TD1* homozygous, none were suppressed by overexpression of *NPY1ΔC*. We observed that overexpression of *NPY1ΔC* actually enhanced the *pid-TD1* phenotypes (Supplemental Figure 7).

### Overexpression of NPY1ΔC disrupts normal responses to auxin and gravity

Overexpression of *NPY1ΔC* caused obvious developmental phenotypes (Figure 3). The *NPY1ΔC OE* plants were smaller with epinastic and darker leaves compared to WT plants (Figure 3A). Overexpression of *NPY1ΔC* led to smaller plant stature and slower development, but *NPY1ΔC OE* plants were eventually able to set seeds (Figure 3B). We noticed that *NPY1ΔC OE* plants often developed small pin-like structures (Figures 3C & 3D). At the seedling stage, *NPY1ΔC* overexpression roots were completely agravitropic (Figure 3E). Moreover, we observed that *NPY1ΔC OE* plants were resistant to auxin (Figure 3F). In media containing 100 nM 2,4- Dichlorophenoxyacetic acid (2,4-D), roots of WT plants stopped growing while roots of *NPY1ΔC OE* plants were able to elongate (Figure 3F). Interestingly, the elongated roots of the *NPY1ΔC OE* plants were still agravitropic (Figure 3F).

**Figure 3.**
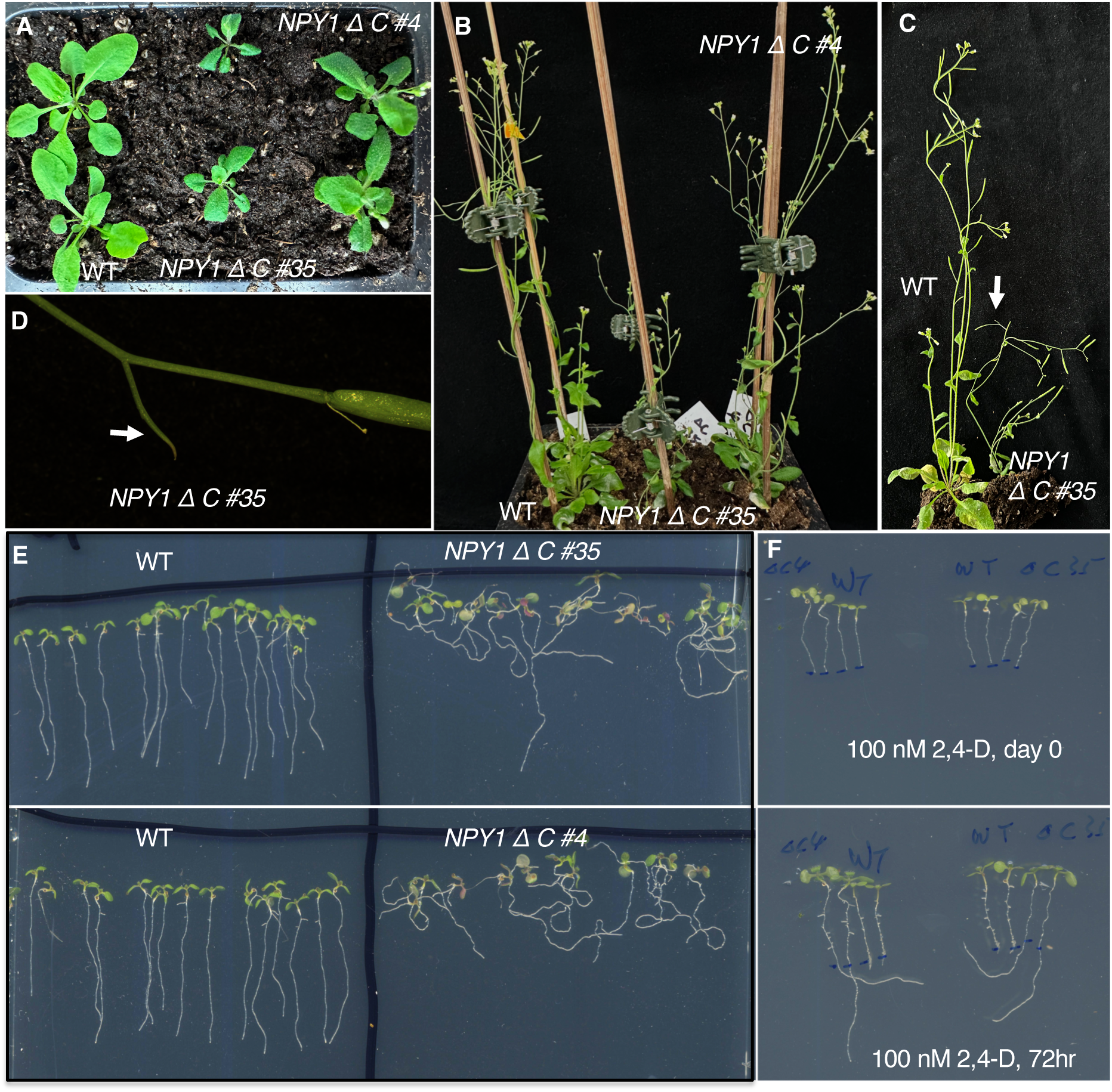
Overexpression of the truncated NPY1 lacking the C-terminal 30 amino acid residues (NPY1ΔC) leads to smaller plants, agravitropic roots, and auxin resistance. A) Overexpression of *NPY1ΔC* leads to small plant stature. The *NPY1ΔC* plants (two independent lines, #35 and #4) have short petiole and epinastic leaves. B) *NPY1ΔC* plants have smaller flowers and are slower in developing siliques. C) *NPY1ΔC* #35 is much shorter than WT. *NPY1ΔC* plants often have small pin-like structures (arrow). D) A pin-like structure (arrow) of *NPY1ΔC* #35 plants. E) *NPY1ΔC* plants have lost normal gravitropic responses. F) NPY1ΔC plants are auxin-resistant. Top panel, 5-day old seedlings were transferred to MS media containing 100 nM 2,4-Dichlorophenoxyacetic acid (2,4-D) and the root tips were marked. Bottom panel: The same plants from the top panel have grown for three more days. Note that roots of WT plants stopped growing, while roots of NPY1ΔC plants continued to grow. 2,4-D did not rescue the defects of gravitropic responses. Plants from left to right: two *NPY1ΔC #4,* two WT, two WT, two *NPY1ΔC #35*.

### Overexpression of NPY1ΔC is sufficient to trigger phosphorylation of PIN proteins

NPY1ΔC protein levels were increased in the analyzed *NPY1ΔC* overexpression lines (#4 and #35) (Figure 4A). The phenotypes of *NPY1ΔC OE* lines appeared to be correlated with the expression levels of NPY1ΔC protein. Line #35 accumulated more NPY1ΔC proteins than line #4 (Figure 4A). Line #35 had stronger phenotypes than line #4 (Figure 3A & B). Overexpression of *NPY1ΔC* had a profound impact on the phospho-proteome (Figure 4 B). In the *NPY1ΔC OE* lines, we detected 18 NPY1 phospho-peptides (Supplemental Table S2, Figure 4C) and the majority of the sites were located in the ANHSPVASVAASSHSPVEK peptide, which was also the main phospho-peptide in the *NPY1 OE* lines (Supplemental Figure 4, Supplemental Table S1).

**Figure 4.**
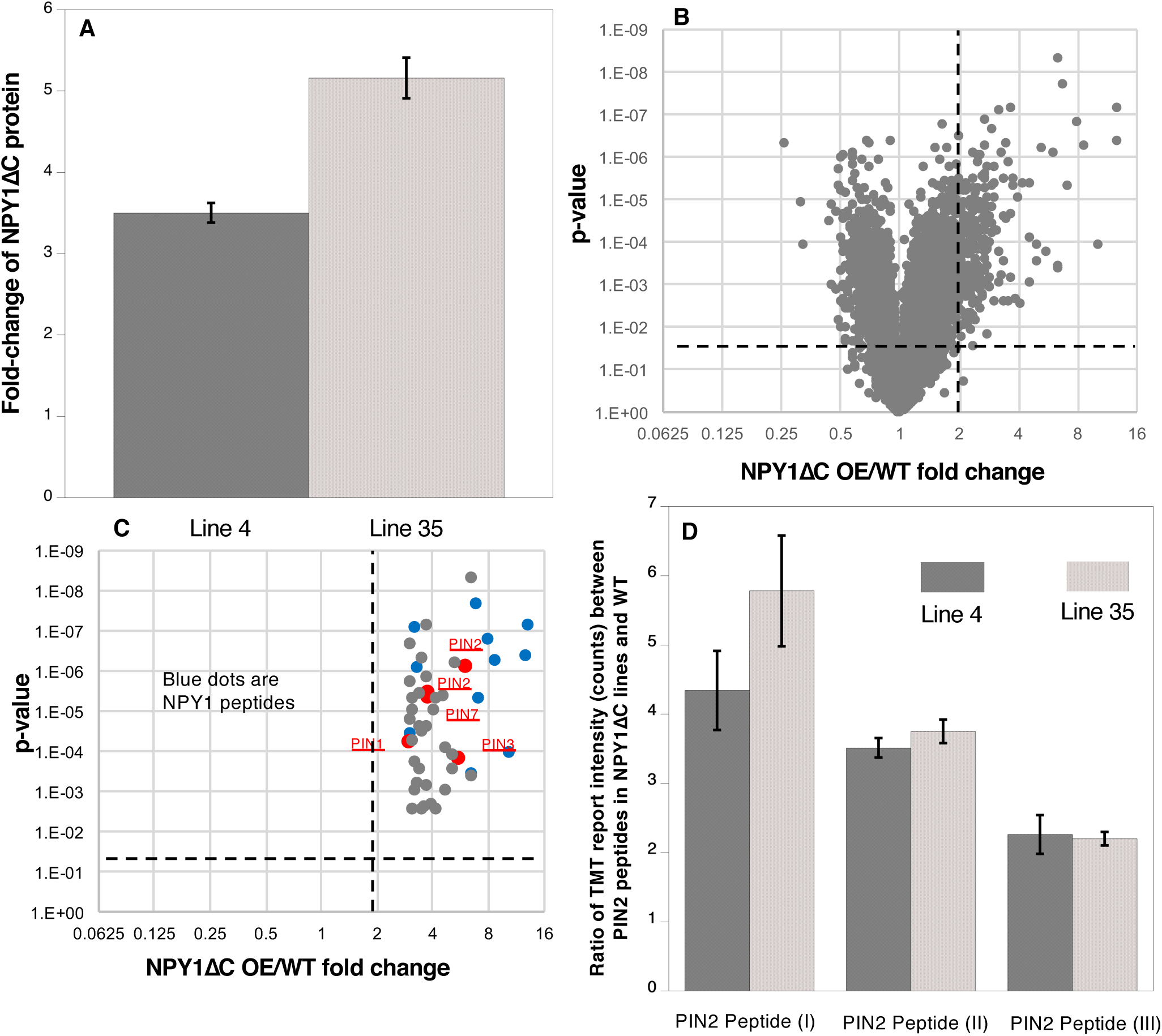
Overexpression of the truncated NPY1 lacking the C-terminal 30 amino acid residues (NPY1ΔC) increases phosphorylation of PIN2 and other PINs. A) NPY1ΔC levels are up several folds in two independent *NPY1ΔC* overexpression lines (Line 4 and Line 35). B) Overexpression of NPY1ΔC led to increases of phosphorylation of many proteins. The phospho-peptides that were at least 2-fold enriched in *NPY1ΔC OE* lines compared to WT and that are statistically significant are shown in the up-right-Quadrant. C) Among the top 50 enriched phospho-peptides in *NPY1 ΔC OE* lines compared to WT, many are from NPY1 (blue dots) and PIN proteins. D) PIN2 phosphorylation increased by at least 2-fold in both line 4 and line 35 for the three detected PIN2 peptides. Peptide I is SESGGGGsGGGVGVGGQNK, peptide II refers to KGsDVEDGGPGPR, and peptide III is HGYTNsYGGAGAGPGGDVYSLQSSK.

Overexpression of *NPY1ΔC* also led to an increase of PIN protein phosphorylation (Figure 4C, Supplemental Table S2). We detected phospho-peptides from all of the long PINs (PIN1, PIN2, PIN3, PIN4, and PIN7) (Supplemental Table S2). We also detected a phospho-peptide from PIN6, which has a shorter hydrophilic loop than the long PINs (Supplemental Table S2).

Many of the identified phosphorylation sites of PIN proteins in the *NPY1ΔC OE* lines were also detected in the *NPY1 OE* lines (Supplemental Figure 8). However, *NPY1ΔC* lines and *NPY1 OE* lines also had their unique phosphorylation sites (Supplemental Figure 8).). A caveat of this comparison is that the phospho-proteomic data were generated using different tissues. We used flowers and inflorescence heads from *NPY1 OE* lines and whole seedings of *NPY1ΔC OE* lines for the analysis. Overall, most of the phosphorylation sites were clustered to regions close to trans-membrane domain 6 (TMD6) (Supplemental Figure 8). Another noticeable feature is that some of the phosphorylated residues were not highly conserved among the PIN proteins (Supplemental Figure 8).

### *NPY1 ΔC*-mediated phosphorylation of PINs correlates with phenotypes similar to those of loss-of-function *pin2* mutants

Very intriguingly, we detected three PIN2 phospho-peptides, which were at least 2-fold more abundant in the *NPY1 ΔC* lines than in WT (Figure 4D, Supplemental Table S2). The level of two of the PIN2 phospho-peptides were higher in the stronger line #35 than in line #4 (Figure 4D). PIN2 is known to play a critical role in gravitropism and *pin2* mutants have agravitropic roots (Chen et al., 1998; Shin et al., 2005). The *NPY1ΔC OE* lines displayed complete agravitropic roots (Figure 3E) and developed small pin inflorescences (Figure 3F), suggesting that *NPY1 ΔC*-mediated phosphorylation of PINs led to their inactivation.

## Discussion

In this paper, we demonstrate that *PID* becomes dispensable for flower initiation when *NPY1* is overexpressed. We further show that overexpression of *NPY1* increases phosphorylation of PIN proteins at sites mostly different from the previously characterized PIN phosphorylation sites. Phosphorylation of PIN proteins at the identified sites described here leads to the inhibition of PIN functions, rather than activation of PINs by phosphorylation at the previously identified sites (Lanassa Bassukas et al., 2022). Moreover, we show that the suppression of *pid* phenotypes by overexpressing *NPY1* requires the C-terminal motif of NPY1. Overexpression of *NPY1ΔC* causes a loss of gravitropism, auxin-resistance in roots, and phosphorylation of PIN proteins. Our results demonstrate that the main function of PID is not to directly phosphorylate PIN proteins and that PID affects the accumulation of NPY1 proteins.

Our findings that overexpression of *NPY1* completely suppressed *pid* mutants demonstrate that *NYP1* functions downstream of *PID* in regulating flower initiation in Arabidopsis (Figure 2). Our data presented in this study are consistent with our previous hypothesis that PID/NPY1-mediated flower initiation uses a mechanism that is analogous to that of phototropism (Figure 5) (Cheng et al., 2007, 2008). The hypothesis was based on the observation that three signaling components in phototropism have their homologous counterparts in the flower initiation pathway (Figure 5). In phototropism, blue light is perceived by phototropins (PHOT1 and PHOT2), which have two LOV domains in the N-terminal part and a kinase domain in the C-terminal part (Huala et al., 1997; Christie et al., 1998; Harper et al., 2003). LOV domains function as photo-receptors that regulate the kinase activities of PHOT1 and PHOT2. The kinase domain of phototropins is highly homologous to PID (Cheng et al., 2007). Downstream of the phototropins is NPH3, which is homologous to NPY1 (Motchoulski and Liscum, 1999; Cheng et al., 2007). Phototropins are the starting point of phototropic signal transduction, and are undoubtedly upstream of NPH3. Our genetic results unambiguously demonstrated that NPY1 functions downstream of PID (Figure 2), further strengthening the hypothesis that phototropism and PID/NPY1 use analogous mechanisms (Figure 5). In dark, NPH3 is phosphorylated and forms a complex with PHOT1 (Pedmale and Liscum, 2007). Upon receiving light, NPH3 is dephosphorylated and transiently dissociated from PHOT1 (Christie et al., 2018; Sullivan et al., 2021; Haga and Sakai, 2023). Continuous light exposure leads to re-constitute PHOT1-NPH3 complex and phosphorylation of NPH3 by PHOT1, suggesting that NPH3 phosphorylation by PHOT1 is rather complicated. We found that NPY1 is phosphorylated regardless of the presence of PID (Figure 2 and Supplemental Table 1). Moreover, we identified a predominant NPY1 phospho-peptide, which is located right after the NPH3 domain (Supplemental Table 1 and Supplemental Figure4). It will be interesting to determine the biological consequences of the phosphorylation sites in NPY1 through mutagenesis. At present, we do not have evidence that PID uses NPY1 as a substrate. However, it is clear that PID has an impact on NPY1 accumulation (Figure 2). Without PID, NPY1 accumulates significantly less (Figure 2).

**Figure 5.**
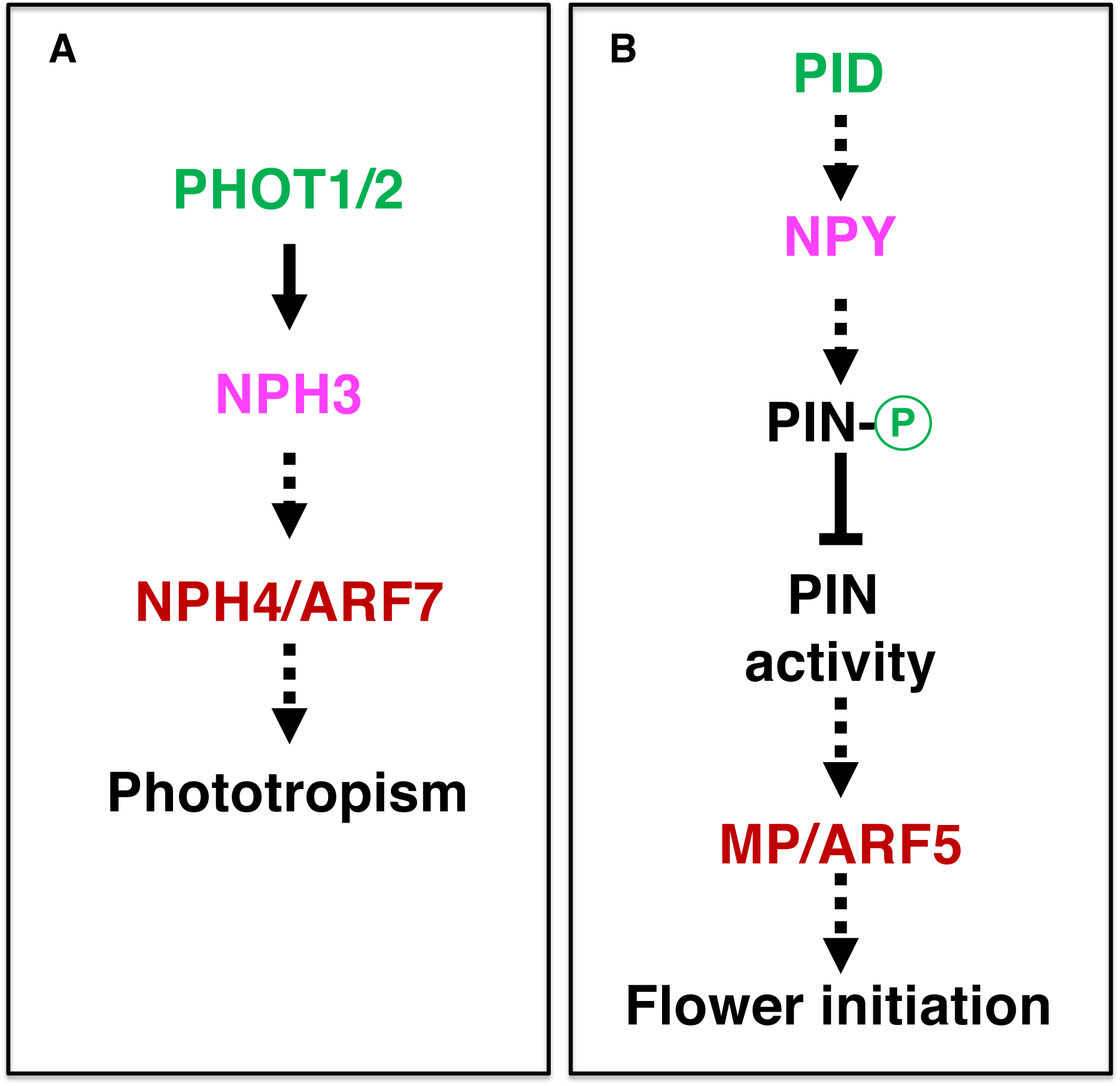
The flower initiation pathway and phototropic pathway use analogous signaling mechanisms. A) Plasma-membrane-localized phototropins perceive blue light, causing changes in phosphorylation status of NPH3 and NPH3-PHOT association. In the nucleus, transcription factor ARF7/NPH4 also plays a role. B) Pathway for auxin-mediated flower initiation. Three genes (*PID*, *NPY*, and *MP*) are homologous to their counterparts in phototropism (color coded). The dotted arrows indicate that there are gaps in our understanding of the step. PHOT: PHOTOTROPIN; NPH3: NON-PHOTOTROPICAL HYPOCOTYL 3; ARF7: AUXIN RESPONSE FACTOR 7; MP: MONOPTEROS.

Unexpectedly, overexpression of *NPY1* leads to phosphorylation of PIN proteins and likely inhibition of PIN functions. The most abundant phospho-peptides in *NPY1 OE* lines are from either NPY1 or PIN proteins (Figure 2). The increased NPY1 protein levels in the *NPY1 OE* lines can partially account for the observed enrichment of phospho-peptides of NPY1 in the overexpression lines. In contrast, overall PIN protein levels were not changed in *NPY1 OE* lines compared to WT, indicating that *NPY1 OE* triggered phosphorylation of PIN proteins. Most of the phosphorylation sites in PIN proteins are not the same as those previously characterized (Lanassa Bassukas et al., 2022). We hypothesize that NPY1-triggered phosphorylation of PINs actually inhibits PIN1 functions, because *NPY1 OE* suppressed *pid* (Figure 2) and because *pid* is suppressed by decreasing PIN1 activity or *PIN1* gene dosage (Mudgett et al., 2023). Our hypothesis is further supported by our results from overexpression of *NPY1ΔC*, which resulted in PIN2 phosphorylation (Figure 4) and agravitropic roots (Figure 3), a phenotype that was observed in *pin2* null mutants (Chen et al., 1998). It is not clear whether *NPY1 OE*-triggered phosphorylation of PINs inhibits auxin transport activity *per se* or disrupts protein-protein interactions between PINs and their partners or both.

Both *NPY1 OE* and *NPY1ΔC* caused great changes in PIN phosphorylation and the phosphorylation led to inhibition of PIN functions (Figures 2 & 4). However, *NPY1 OE* suppressed *pid* mutants whereas *NPY1ΔC* could not suppress *pid* (Figure 2 & 3), suggesting that inhibition of PIN functions *per se* is not sufficient for suppression of *pid*. It is known that PIN and NPY proteins physically interact to form protein complexes (Glanc et al., 2021; Matthes et al., 2024). Our results suggest that the C-terminal domain of NPY1 likely interacts with an unknown protein, which is required for normal PID and PIN functions and which needs to be recruited to the NPY-PIN complexes.

In previous models regarding the mechanisms of PID, PIN1, and NPY1, the first step is direct phosphorylation of PINs by PID (Friml et al., 2004; Michniewicz et al., 2007). Phosphorylated PINs recruit NPY1 to the plasma membrane to maintain PIN polarity by limiting lateral diffusion of the PIN complex (Glanc et al., 2021). However, that overexpression of *NPY1* eliminates the need of *PID* in flower initiation and that overexpression of *NPY1* leads to phosphorylation of PIN proteins in the absence of PID are not consistent with the model that PIN1 needs to be directly phosphorylated by PID to be activated. The fact that *pin1* and *pid* displayed opposite cotyledon/leaf phenotypes (Bennett et al., 1995) and that *pid* is suppressed by heterozygous *pin1* mutants (Figure 1) and PIN1-GFPHDR (Mudgett et al., 2023) demonstrated that PID and PIN1 function in opposite directions. Our results firmly established the relative positions of PID and NPY1in the pathway responsible for flower initiation in Arabidopsis (Figure 5). Moreover, we show that PIN1 functions downstream of NPY1, which recruits unknown kinase(s) to phosphorylate PIN1 and to inhibit PIN1 (Figure 5).

## Materials and Methods

### Plant Growth

Plants used in this study were the *Arabidopsis thaliana* Columbia-0 ecotype. Sterilized seeds were sown on a 0.7% agar-agar medium containing Murashige and Skoog basal salts at ½ concentration. Sown seeds were subjected to a two-day dark stratification period at 4°C and then moved to a growth chamber with a 16-hour day / 8-hour night cycle. After one week, seedlings were transferred to soil and maintained under the same light conditions.

### Plant Transformation

Transgenic plants were created by performing the floral dipping procedure on adult plants with unopened flowers using published protocols (Clough and Bent, 1998). The *Agrobacterium tumefaciens* strain GV3101 was used to perform all transformations in this study. Seeds were harvested from transformed plants and transformants were selected via the expression of the mCherry fluorescent marker present on the transformed vectors (Gao et al., 2016).

### Plasmid Construction

Vectors for gene knockouts were created by cloning two guide RNAs into the pHEE401E backbone as described in (Mudgett et al., 2024). Overexpression vectors were assembled via Gibson Assembly in the pHDE backbone (Gao et al., 2016). Primers used in this study are listed in the Supplemental Table 3.

### Proteomics Method

The following tissues were collected and were immediately frozen in liquid nitrogen for proteomic analysis. Inflorescence heads, which include flower buds and meristems from WT (Columbia background)-68, *NPY1 OE* in WT-68, *NPY1 OE* in *pid-c1*-68, WT-83, *NPY1 OE* in WT-83, *NPY1 OE* in *pid-c1*-83 were collected. Whole seedlings of 7-day old Col-WT and *NPY1ΔC* #35, and #4 were used in this study.

About 0.5 gram of frozen tissue was ground in liquid nitrogen by a mortar/pestle for 15 minutes to fine powders, and then transferred to a 50ml conical tube. Proteins were precipitated and washed by 50 ml -20 °C acetone three times, then by 50 ml -20 °C methanol three times. Samples were centrifuged at 4,000x g, 4 °C for 10 minutes. Supernatant was removed and discarded.

Protein pellets were suspended in extraction buffer (8 M Urea/100mM Tris/5mM TCEP/phosphatase inhibitors, pH 7). Proteins were first digested with Lys-C (Wako Chemicals, 125-05061) at 37 °C for 15 minutes. Protein solution was diluted 8 times to 1M urea with 100mM Tris and digested with trypsin (Roche, 03708969001) for 12 hours. Cysteines were alkylated by adding 10 mM iodoacetamode and incubating at 37 °C for 30 minutes in dark.

Digested peptides were purified on a Waters Sep-Pak C18 cartridges, eluted with 60% acetonitrile. TMT labeling (Supplemental Tabel S4) was performed in 60% acetonitrile/100mM Hepes, pH 7. TMT labeling efficiency was checked by LC-MS/MS to be greater than 99%. Labeled peptides from different samples were pooled together. 150 µg of pooled peptides were analyzed by 2D-nano-LC-MS/MS for total proteome profiling and 1 mg of total peptides was used for phosphopeptide enrichment.

Phosphopeptide enrichment was performed using CeO2 affinity capture. 20% colloidal CeO2 (Sigma, 289744) was added to the acidified peptide solution (2% TFA/2M lactic acid/60% acetonitrile). After brief vortexing, CeO2 with captured phosphopeptides was spun down at 5,000 g for 1 minute. Supernatant was then removed and the CeO2 pellet was washed with 1 mL of 2% TFA/2M lactic acid/50% acetonitrile. Phosphopeptides were eluted by adding 200 µL eluting buffer (50mM (NH4)2HPO4, 2M NH3.H2O, 10mM EDTA, pH 9.5) and vortexed briefly. CeO2 was precipitated by adding 200 µL acetonitrile. Samples were centrifuged at 16,100 g for 1 minute. Supernatant containing phosphopeptides was removed and dried in a Speedvac. Phosphopeptides were resuspended in 100 mM citric acid and ready for mass spec analysis.

A Thermo Scientific Vanquish Neo UHPLC system (Buffer A: Water with 0.1% formic acid; Buffer B: 80% acetonitrile with 0.1% formic acid) was used to deliver a flow rate of 500 nL/min to a self-packed 3-phase capillary (360 µm OD/200 µm ID) chromatography column. Column phases were a 10 cm long reverse phase (RP1, 5 µm Zorbax SB-C18, Agilent), 6 cm long strong cation exchange (SCX, 3 µm PolySulfoethyl, PolyLC), and 20 cm long reverse phase 2 (RP2, ReproSil-Pur 120 C18-AQ, 1.9 µm), with the electrospray tip of the fused silica tubing pulled to a sharp tip.

Peptide mixtures were loaded onto RP1 using an off-line pressure chamber; and the 3 sections were joined for on-line 2D LC separation. Peptides were eluted from RP1 section to SCX section using a 0 to 80% acetonitrile gradient for 60 minutes, and then were fractionated by the SCX column section by injecting a series of ammonium acetate solutions (5 µL of 10 mM, 20 mM, 30 mM, 40 mM, 50 mM, 60 mM, 70 mM, 80 mM, 90 mM, 100 mM, and 1 M), followed by high-resolution reverse phase separation on the RP2 section of the column using an acetonitrile gradient (0–0.1 min 1% B to 5% B, 0.1–110.1 min 5% B to 35% B, 110.1–120.1 min 35% B to 60% B, 120.1–120.3 min 60% B to 95% B, 120.3–126 min 95% B).

Mass Spectra were acquired on a Thermo Exploris 480 mass spectrometer operated in positive ion mode with a source temperature of 300 °C and spray voltage of 2.1 kV. Automated data-dependent acquisition was employed of the top 20 ions with an isolation window of 0.7 Da and collision energy of 35. Precursor Fit was set to 70% of fit threshold and 0.7 Da fit window. The mass resolution is set at 100,000 for MS and 30,000 for MS/MS scans, respectively. TurboTMT was enabled. Dynamic exclusion of 30 seconds was used to improve the duty cycle.

The raw data was extracted and searched using Spectrum Mill vBI.07 (Broad Institute of MIT and Harvard). MS/MS spectra with a sequence tag length of 1 or less were considered to be poor spectra and were discarded. The remaining high-quality MS/MS spectra were searched against Arabidopsis TAIR11 protein database. A 1:1 concatenated forward-reverse database was constructed to calculate the false discovery rate (FDR). Common contaminants such as trypsin and keratin were included in the protein database. There were 96,562 protein sequences in the final protein database. Search parameters were set to Spectrum Mill’s default settings with the enzyme parameter limited to full tryptic peptides with a maximum mis-cleavage of 1. Cutoff scores were dynamically assigned to each dataset to obtain the false discovery rates (FDR) of 0.1% for peptides, and 1% for proteins. Phosphorylation sites were localized to a particular amino acid within a peptide using the variable modification localization (VML) score in Spectrum Mill software. Proteins that share common peptides were grouped using principles of parsimony to address protein database redundancy. Total TMT reporter intensities were used for relative protein quantitation. Peptides shared among different protein groups were removed before TMT quantitation. Isotope impurities of TMT reagents were corrected using correction factors provided by the manufacturer (Thermo). Median normalization was performed to normalize the protein TMT reporter intensities in which the log ratios between different TMT tags were adjusted globally such that the median log ratio was zero.

### Proteomics data deposition

The raw spectra for the proteome data have been deposited in the Mass Spectrometry Interactive Virtual Environment (MassIVE) repository (massive.ucsd.edu/ProteoSAFe/static/massive.jsp, accession ID MSV000098241). FTP download link before publication: ftp://MSV000098241@massive-ftp.ucsd.edu; FTP download link after publication: ftp://massive-ftp.ucsd.edu/v10/MSV000098241/.

## Acknowledgments

We thank the UCSD Goeddel Family Technology Sandbox for providing the Thermo Scientific Vanquish Neo UHPLC system and Exploris 480 mass spectrometer used in this research to generate the proteomics and phospho-proteomics data. This work was partially supported by the NIH Cell and Molecular Genetics Training Program (5T32GM007240-43) to M. M., NSF Grant 1546899 to S.P.B., and Tata Chancellor’s Endowed Professorship to Y. Z.

## Legends for Supplemental Figures

**Supplemental Figure S1. Genetic interactions between *pid* and *npy* mutants.** A) The phenotypes of *pid-TD1* is not rescued by heterozygous *npy1-2*, which is a T-DNA insertion null allele. B) The *npy1-2* mutant is enhanced by heterozygous *pid-TD1* mutant. Plants with *npy1-2 pid-TD1^+/-^* genotype often make pin-like inflorescences (arrow)

**Supplemental Figure S2. Overexpression of *NPY1* rescues *pid* mutants and causes obvious developmental phenotypes.** A) Young adult *NPY1 OE* plants in WT and in *pid-c1*. The WT control was at the left. Note that the *NPY1OE* plants had short petioles in both WT and in *pid-c1* backgrounds. B) *NPY1 OE* plants had more compact inflorescences and shorter stature compared to WT. C) *NPY1 OE* completely eliminated pin-structures in *pid-c1* and developed normal looking inflorescences head. D) Dissected flowers of WT and *NPY1 OE* plants. Note that flowers of *NPY1 OE* in *pid-c1* occasionally have minor defects such as fewer stamens and fused organ (arrow).

**Supplemental Figure S3. Overexpression of *NPY1* is sufficient to rescue *pid* null mutants.** A) Overexpression of *NPY1* rescues *pid-TD1*, which harbors a T-DNA insertion in the second exon and which is a strong allele. B) Another T-DNA *pid* mutant, which also has the T-DNA insertion in the second exon, is rescued by overexpression of *NPY1*. C) A close-up of the panel B. Some of the rescued *pid-TD2* siliques only have one valve (arrows).

**Supplemental Figure S4. The identified phosphorylation sites in *NPY1*.** The NPY protein sequences along with NPH3 protein were aligned using ClustalW. Top panel shows the phosphorylation site S181 in NPY1. S181 is located in a highly conserved WSYT motif and is located in the region that separates the BTB domain and NPH3 domain. Middle panel shows a phosphorylation site inside the NPH3 domain. The bottom panel shows the predominant phospho-peptide ANHSPVASVAASSHSPVEK, which is located immediately after the NPH3 domain. Although the serine residues appear not conserved among NPY proteins, this region is rich in Ser/Thr residues in other NPY proteins and in NPH3. Previous studies have identified two phosphorylation sites in NPY1: S514 and S553, which are marked green. S553 was also identified as a phosphorylation site in this study (underlined green). S514 containing peptide was identified in this study, but it was S523 that was phosphorylated. The C-terminal “SIS” motif in NPY1 also exists in NPH3. Previous studies have shown that the two serine residues in “SIS” (in bold) in NPH3 are phosphorylated

**Supplemental Figure S5. Overexpression of *NPY1* leads to increases of phosphorylation of PIN proteins.** The hydrophilic loops of the “long PINs” (PIN1,2,3,4,7) were aligned using ClustalW. The previously characterized phosphorylation sites are in bold black. The phosphorylated residues identified in *NPY1* overexpression (OE) lines are highlighted in bold red. In the *NPY1* overexpression lines, 10 phospho-peptides of PIN3 were identified and 9 of them were significantly increased in the *NPY1 OE* lines compared to WT. Seven PIN7 phospho-peptides were identified, but only the peptide with S238 was significantly increased in *NPY1 OE* lines. More than 4-fold increase in phosphorylation at S405 of PIN4 in *NPY1 OE* lines was observed. Phosphorylation of two highly conserved serine residues in PIN1 was increased in the NPY1 overexpression lines.

**Supplemental Figure S6. Potential roles of the C-terminal domain of NPY1**. A) The C-terminal domain (104 amino acid residues), which is located immediately after the conserved NPH3 domain, is rich in Ser/Thr (in bold). The C-terminal fragment (30 amino acid residues) contains 50% of Ser. The NPY1ΔC construct deleted the C-terminal 30 amino acid residues (highlighted in red). B) Potential PID-NPY1 interaction predicted from Alphafold3. The PID (gold)-NPY1(blue)-ATP multimer predicted by the Alphafold3 webserver is shown. The disordered C-terminal tail of NPY1 was predicted to associate with the PID active site (red box). (C) Close-up of the predicted association between the NPY1 C-terminal tail and the PID kinase active site. Note the proximity of ATP, the catalytic aspartate D205 of PID, and S569, a possible phosphorylation target in NPY1 (see panel A). NPH3 contains a similar C-terminal motif to NPY1, and phosphorylation of the S569 counterpart in NPH3 (S744) by PHOT1 was shown to be involved in light signaling. (https://doi.org/10.1038/s41467-021-26333-5)

**Supplemental Figure S7. Overexpression of *NPY1ΔC* was not able to suppress *pid-TD1*.** Overexpression of *NPY1ΔC* in *pid-TD1* resulted in a smaller plant with pin-like inflorescences

**Supplemental Figure S8. Overexpression of NPY1 lacking the C-terminal 30 amino acid residues (*NPY1ΔC*) leads to increases of phosphorylation of PIN proteins.** The hydrophilic loops of the “long PINs” (PIN1,2,3,4,7) are shown. The previously characterized phosphorylation sites are in bold & black. The phosphorylated residues identified in *NPY1* OE lines, but not identified in the *NPY1ΔC OE* lines are highlighted in bold & yellow. The phosphorylated residues identified only in the *NPY1ΔC OE* lines are highlighted in bold & green. Residues highlighted in purple & bold are identified in both *NPY1 OE* lines and the *NPY1ΔC* lines.

**Supplemental Table S1. Phospho-peptides from NPY1 protein in both WT and *pid-c1* backgrounds**. *NPY1* overexpression T1 plants with *pid-c1*^+/-^ genotype was self-pollinated. At T2 stage, three genotypes were selected: WT without transgenes and without *pid-c1* mutation (this is called WT-xx, xx refers to the line number); NPY1 OE without *pid-c1* mutation (Called NPY1 in WT xx); NPY1 OE in *pid c1* (called NPY1 in *pid* xx). Inflorescence heads with flower buds of the lines were used for proteomic analysis. All of the detected NPY1peptides were more abundant in the overexpression lines than in WT. Two most abundant peptides in NPY1 OE lines were highlighted yellow. PID did not affect NPY1 phosphorylation, as all of the NPY1 phospho-peptides were detected in *pid-c1* background.

**Supplemental Table S2. Phosphorylation of NPY1 and PIN proteins caused by *NPY1ΔC* overexpression**. L4 and L35 refer to *NPY1ΔC OE* line #4 and #35, respectively. WT: wild type. Lower case s or t in the peptides refers to the phosphorylated residues. The PIN2 phospho-peptides are highlighted in yellow, which were enriched in the *NPY1ΔC OE* lines.

**Supplemental Table S3. Primers used in this study.**

**Supplemental Table S4. TMT labelling schemes**

## References

Barbosa IC, Zourelidou M, Willige BC, Weller B, Schwechheimer C (2014) D6 PROTEIN KINASE activates auxin transport-dependent growth and PIN-FORMED phosphorylation at the plasma membrane. Dev Cell 29: 674–685

Barbosa ICR, Hammes UZ, Schwechheimer C (2018) Activation and Polarity Control of PIN-FORMED Auxin Transporters by Phosphorylation. Trends Plant Sci 23: 523–538

Bennett SRM, Alvarez J, Bossinger G, Smyth DR (1995) Morphogenesis in pinoid mutants of Arabidopsis thaliana. The Plant Journal 8: 505–520

Chen R, Hilson P, Sedbrook J, Rosen E, Caspar T, Masson PH (1998) The arabidopsis thaliana AGRAVITROPIC 1 gene encodes a component of the polar-auxin-transport efflux carrier. Proc Natl Acad Sci U S A 95: 15112–15117

Cheng Y, Dai X, Zhao Y (2006) Auxin biosynthesis by the YUCCA flavin monooxygenases controls the formation of floral organs and vascular tissues in Arabidopsis. Genes Dev 20: 1790–1799

Cheng Y, Dai X, Zhao Y (2007) Auxin synthesized by the YUCCA flavin monooxygenases is essential for embryogenesis and leaf formation in Arabidopsis. Plant Cell 19: 2430–2439

Cheng Y, Qin G, Dai X, Zhao Y (2007) NPY1, a BTB-NPH3-like protein, plays a critical role in auxin-regulated organogenesis in Arabidopsis. Proc Natl Acad Sci U S A 104: 18825–18829

Cheng Y, Qin G, Dai X, Zhao Y (2008) NPY genes and AGC kinases define two key steps in auxin-mediated organogenesis in Arabidopsis. Proc Natl Acad Sci U S A 105: 21017–21022

Cheng Y, Zhao Y (2007) A Role for Auxin in Flower Development. Journal of Integrative Plant Biology 49: 99–104

Christensen SK, Dagenais N, Chory J, Weigel D (2000) Regulation of auxin response by the protein kinase PINOID. Cell 100: 469–478

Christie JM, Reymond P, Powell GK, Bernasconi P, Raibekas AA, Liscum E, Briggs WR (1998) Arabidopsis NPH1: a flavoprotein with the properties of a photoreceptor for phototropism. Science 282: 1698–1701

Christie JM, Suetsugu N, Sullivan S, Wada M (2018) Shining Light on the Function of NPH3/RPT2-Like Proteins in Phototropin Signaling. Plant Physiol 176: 1015–1024

Clough SJ, Bent AF (1998) Floral dip: a simplified method for Agrobacterium-mediated transformation of Arabidopsis thaliana. Plant J 16: 735–743

Friml J, Yang X, Michniewicz M, Weijers D, Quint A, Tietz O, Benjamins R, Ouwerkerk PB, Ljung K, Sandberg G, Hooykaas PJ, Palme K, Offringa R (2004) A PINOID-dependent binary switch in apical-basal PIN polar targeting directs auxin efflux. Science 306: 862–865

Furutani M, Kajiwara T, Kato T, Treml BS, Stockum C, Torres-Ruiz RA, Tasaka M (2007) The gene MACCHI-BOU 4/ENHANCER OF PINOID encodes a NPH3-like protein and reveals similarities between organogenesis and phototropism at the molecular level. Development 134: 3849–3859

Furutani M, Sakamoto N, Yoshida S, Kajiwara T, Robert HS, Friml J, Tasaka M (2011) Polar-localized NPH3-like proteins regulate polarity and endocytosis of PIN-FORMED auxin efflux carriers. Development 138: 2069–2078

Furutani M, Vernoux T, Traas J, Kato T, Tasaka M, Aida M (2004) PIN-FORMED1 and PINOID regulate boundary formation and cotyledon development in Arabidopsis embryogenesis. Development 131: 5021–5030

Galweiler L, Guan C, Muller A, Wisman E, Mendgen K, Yephremov A, Palme K (1998) Regulation of polar auxin transport by AtPIN1 in Arabidopsis vascular tissue. Science 282: 2226–2230

Gao X, Chen J, Dai X, Zhang D, Zhao Y (2016) An Effective Strategy for Reliably Isolating Heritable and Cas9-Free Arabidopsis Mutants Generated by CRISPR/Cas9-Mediated Genome Editing. Plant Physiol 171: 1794–1800

Glanc M, Van Gelderen K, Hoermayer L, Tan S, Naramoto S, Zhang X, Domjan D, Vcelarova L, Hauschild R, Johnson A, de Koning E, van Dop M, Rademacher E, Janson S, Wei X, Molnar G, Fendrych M, De Rybel B, Offringa R, Friml J (2021) AGC kinases and MAB4/MEL proteins maintain PIN polarity by limiting lateral diffusion in plant cells. Curr Biol 31: 1918–1930 e1915

Haga K, Sakai T (2023) Photosensory adaptation mechanisms in hypocotyl phototropism: how plants recognize the direction of a light source. J Exp Bot 74: 1758–1769

Harper SM, Neil LC, Gardner KH (2003) Structural basis of a phototropin light switch. Science 301: 1541–1544

Huala E, Oeller PW, Liscum E, Han IS, Larsen E, Briggs WR (1997) Arabidopsis NPH1: a protein kinase with a putative redox-sensing domain. Science 278: 2120–2123

Lanassa Bassukas AE, Xiao Y, Schwechheimer C (2022) Phosphorylation control of PIN auxin transporters. Curr Opin Plant Biol 65: 102146

Matthes MS, Yun N, Luichtl M, Büschges U, Fiesselmann BS, Strickland B, Lehnardt MS, Ruiz RAT (2024) Separate domains of the *Arabidopsis* ENHANCER OF PINOID drive its own polarization and recruit PIN1 to the plasma membrane. bioRxiv: 2024.2003.2011.584374

Michniewicz M, Zago MK, Abas L, Weijers D, Schweighofer A, Meskiene I, Heisler MG, Ohno C, Zhang J, Huang F, Schwab R, Weigel D, Meyerowitz EM, Luschnig C, Offringa R, Friml J (2007) Antagonistic regulation of PIN phosphorylation by PP2A and PINOID directs auxin flux. Cell 130: 1044–1056

Motchoulski A, Liscum E (1999) Arabidopsis NPH3: A NPH1 photoreceptor-interacting protein essential for phototropism. Science 286: 961–964

Mudgett M, Abramson B, Dai X, Kang R, Young E, Michael T, Zhao Y (2024) Letter to the Editor: Gene Targeting in Arabidopsis through One-Armed Homology-Directed Repair. Plant Cell Physiol 65: 1937–1940

Mudgett M, Shen Z, Dai X, Briggs SP, Zhao Y (2023) Suppression of pinoid mutant phenotypes by mutations in PIN-FORMED 1 and PIN1-GFP fusion. Proc Natl Acad Sci U S A 120: e2312918120

Okada K, Ueda J, Komaki MK, Bell CJ, Shimura Y (1991) Requirement of the Auxin Polar Transport System in Early Stages of Arabidopsis Floral Bud Formation. Plant Cell 3: 677–684

Pedmale UV, Liscum E (2007) Regulation of phototropic signaling in Arabidopsis via phosphorylation state changes in the phototropin 1-interacting protein NPH3. J Biol Chem 282: 19992–20001

Przemeck GK, Mattsson J, Hardtke CS, Sung ZR, Berleth T (1996) Studies on the role of the Arabidopsis gene MONOPTEROS in vascular development and plant cell axialization. Planta 200: 229–237

Shin H, Shin HS, Guo Z, Blancaflor EB, Masson PH, Chen R (2005) Complex regulation of Arabidopsis AGR1/PIN2-mediated root gravitropic response and basipetal auxin transport by cantharidin-sensitive protein phosphatases. Plant J 42: 188–200

Smyth D, Alvarez J (1995) Morphogenesis in pinoid mutants of Arabidopsis thaliana. Plant Journal 8: 505–520

Sullivan S, Waksman T, Paliogianni D, Henderson L, Lutkemeyer M, Suetsugu N, Christie JM (2021) Regulation of plant phototropic growth by NPH3/RPT2-like substrate phosphorylation and 14-3-3 binding. Nat Commun 12: 6129

Treml BS, Winderl S, Radykewicz R, Herz M, Schweizer G, Hutzler P, Glawischnig E, Ruiz RA (2005) The gene ENHANCER OF PINOID controls cotyledon development in the Arabidopsis embryo. Development 132: 4063–4074

Willige BC, Ahlers S, Zourelidou M, Barbosa IC, Demarsy E, Trevisan M, Davis PA, Roelfsema MR, Hangarter R, Fankhauser C, Schwechheimer C (2013) D6PK AGCVIII kinases are required for auxin transport and phototropic hypocotyl bending in Arabidopsis. Plant Cell 25: 1674–1688

Yamaguchi N, Wu MF, Winter CM, Berns MC, Nole-Wilson S, Yamaguchi A, Coupland G, Krizek BA, Wagner D (2013) A molecular framework for auxin-mediated initiation of flower primordia. Dev Cell 24: 271–282

Yang Z, Xia J, Hong J, Zhang C, Wei H, Ying W, Sun C, Sun L, Mao Y, Gao Y, Tan S, Friml J, Li D, Liu X, Sun L (2022) Structural insights into auxin recognition and efflux by Arabidopsis PIN1. Nature 609: 611–615

